# The olivocerebellar system differentially encodes the effect sensory events exert on behavior

**DOI:** 10.1101/2025.01.14.632916

**Authors:** Irina Scheer, Mario Prsa

## Abstract

Inferior olive neurons convey information about sensorimotor events via climbing fibers to the cerebellum, but their functional significance remains unclear. We directly imaged, with two-photon microscopy, climbing fiber axonal terminals in the cerebellum during a task that successively exposed mice to a force perturbation, a movement instruction and reward; each followed by multiple modes of motor activity. Climbing fiber activations by the sensory events were either generic or informative about the consequences the encoded event has on behavior. The number of informative cells and the information strength are regulated by event modality and functional complexity of the cell’s activity. We observed an additional, previously unreported activation of climbing fibers: they carried probabilistic information on the behavioral context during idle waiting periods preceding stimulus presentation. Our findings reveal properties of olivary neurons that are key for defining their function in the cerebellum-dependent control of behavior. They suggest that the inferior olive flexibly instructs the cerebellum of any process that may shape an animal’s action.

## Introduction

The cerebellum is a powerful motor coordination center. It receives rich sensory, motor and contextual information via mossy fiber inputs from the sensory periphery, neocortical motor and cognitive areas and cerebellar nucleo-cortical feedback ^1–5^. The cerebellum selectively sorts out this extensive information within a rigorously structured internal circuit and funnels pertinent signals via a limited number of nuclear efferent cells ^6^ to enable efficient behavioral execution. The key pruning process occurs at the dendritic tree of Purkinje cells. Thousands of parallel fiber synapses, relaying mossy fiber inputs to Purkinje cells, are modified throughout the entire dendritic tree, by an instructive signal from a single climbing fiber (CF) originating in the inferior olive ^7^.

Early theoretical work proposed that CFs provide information about movements and movement errors to the cerebellum for correcting and adapting motor performance ^8,9^. This has remained arguably the most influential hypothesis about the function of the olivocerebellar system. In the context of forelimb reaching, it might indeed seem that olivary neurons primarily encode movement kinematics and performance errors ^10–17^. However, by extending the types of behaviors studied, in-vivo recordings reveal that CFs carry a diversity of signals to the cerebellum. The most effective stimuli seem to be unexpected tactile or proprioceptive perturbations of a particular body part, even under anesthesia ^18–20^. This is consistent with the inferior olive receiving abundant afferents from spinal cord, trigeminal and dorsal column nuclei and maintaining a well-defined somatotopy of the contralateral body ^21–24^. Olivary neurons therefore broadly respond to sensory disturbances rather than movement errors per se ^25–27^ and can show no, little or poorly time locked activation during active movements ^27–29^. Furthermore, remote “teleceptive” stimuli with no physical contact with the animal (e.g. neutral visual or auditory cues) also evoke reliable olivary responses ^25,26,30^, multiple sensory modalities and behavioral variables can be represented by individual cells ^25,26,28,31,32^ and CF inputs to forelimb areas of the cerebellum are more prominently modulated by reward related signals than movements ^33,34^. Such multifaceted information is likely mediated by the combination of direct sensory pathways with cerebral inputs (via the mesodiencephalic junction) to the inferior olive ^35,36^.

It therefore seems that the inferior olive reports the occurrence of generic sensory events rather than stimuli with specific properties (e.g. movement errors). What function might this seemingly generic olivary activation accomplish? One hypothesis is that CF activity is best understood as a signal about stimulus saliency allowing the animal to organize the transition between successive behavioral states or contexts ^25,26,32,37^. For example, CF bursts time locked to reward onset might act as a trigger to switch behavior between reward anticipation and its consumption ^33^. It might in this manner attach meaning to any arbitrary sensory stimulus that has, through associative learning, gained behavioral relevance^32^. Alternatively, or complementarily, it might be predictive of upcoming motor behavior ^16,31,38^ and thereby dictate how to organize movement parameters in response to a sensory stimulus. The reward response would in that case be modulated by lick latency, rate or precision ^38^. We refer to these two as the “sensory saliency” and “motor organization” hypotheses, respectively. “Sensory saliency” implies that CF activation is generic in nature; it is triggered by task relevant cues irrespective of how the subsequent behavioral transition is made. “Motor organization” on the other hand implies that CF activation is instructive in nature; it has an influence on behavioral execution and is thereby expected to be informative about the initiation or kinematics of ensuing movements.

To understand the information signaled by the olivocerebellar system, we here imaged climbing fibers in a task comprising successive sensory events, each with a different behavioral significance. It allowed us to evaluate the interaction between CF responses to forelimb force perturbations, instructive “teleceptive” cues or reward, and subsequent forelimb movement or licking behavior. We observe that “sensory saliency” and “motor organization” labels can be attributed to CFs depending on the modality that produces the response as well as its functional complexity. Moreover, our results indicate that many CFs are modulated during silent waiting periods based on the nature of the anticipated sensory event.

## Results

### Partitioned representation of multimodal cues by cerebellar climbing fibers

We designed a task in which mice responded to three successive sensory events, each having a different behavioral significance. Head-fixed mice were trained to grasp and hold the endpoint of a single-axis robotic manipulandum (Fig. 1A). At the beginning of each trial, the endpoint was held at the starting home position. After a period of continuous holding, a resistive force was applied by the actuating motor. At force onset, the limb position was either perturbed (posterior displacement; red arrow Fig. 1B) or remained stable as the mice produced sufficient force to counteract the perturbation (adapted, green trace, Fig. 1B). After a fixed delay, an auditory go cue instructed the mice to initiate a forward movement towards a target position. Perturbed positions resulted in poorly timed and slow movements (red trace, Fig. 1B) whereas adapted movements were either cued (initiated after the go cue; green trace, Fig. 1B), uncued (initiated prior to the go cue) or no movement was produced (black traces, Fig. 1B). After reaching the target, the manipulandum was held at position for a fixed delay and a reward was delivered. Trials with releases of the manipulandum at any time after force onset or with movements reaching the target prior to go cue were aborted (see Methods for details).

**Figure 1.**
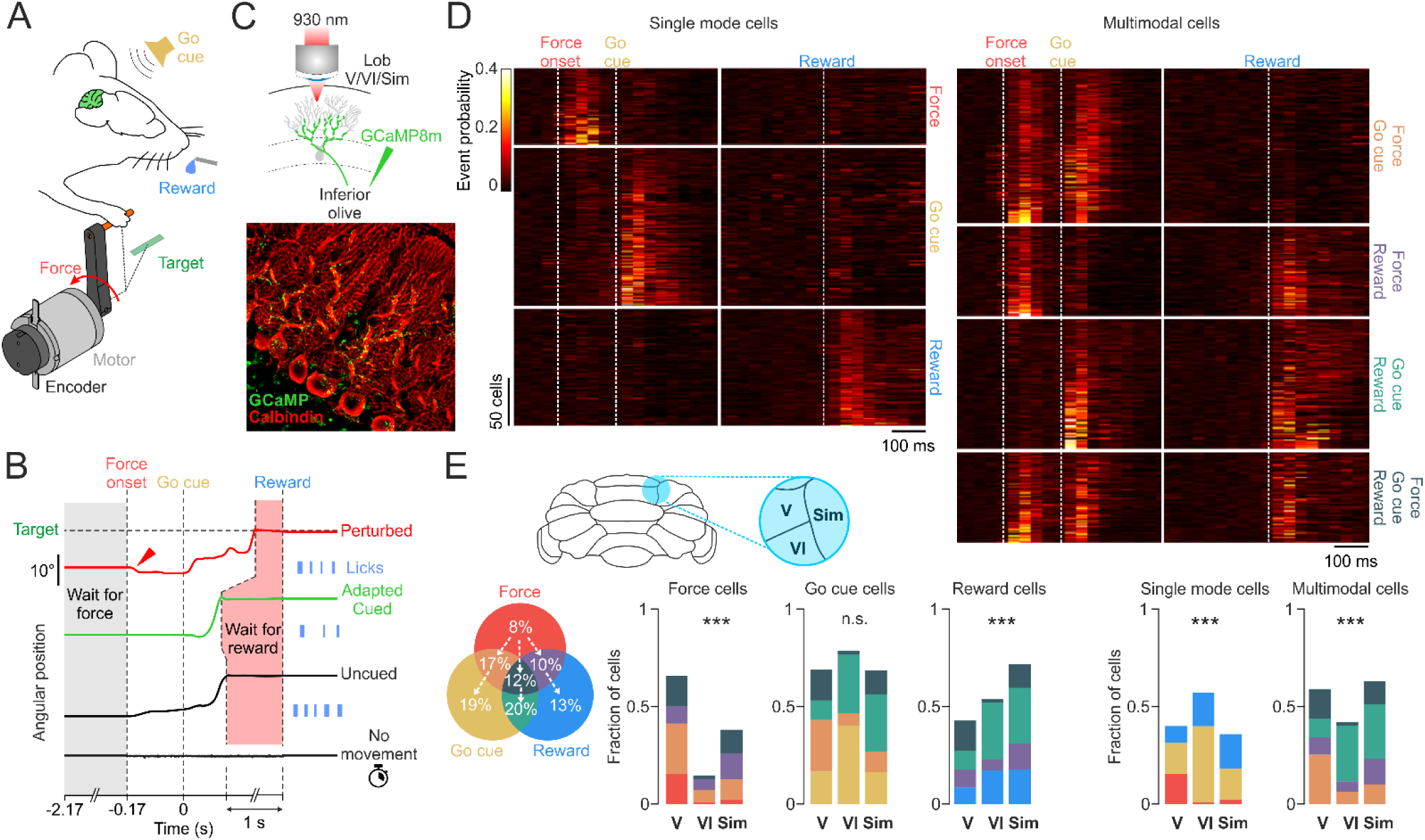
Climbing fibers exhibit pure and mixed selectivity unevenly across lobules V, VI and Simplex. **A:** Mice were trained to execute cued forelimb movements with a robotic manipulandum to obtain rewards following a force perturbation. **B:** Task timeline showing four trial types: limb position perturbed by the force (red arrow) followed by a slow movement to target (red trace), limb position adapted to the force followed by a correctly timed movement to target (green trace), movement initiation preceding the go cue (top black trace) and absence of movement (bottom black trace). The shaded areas are silent periods of continuous holding of the manipulandum without movements and sensory cues defined as “wait for force” (grey) and “wait for reward” (red). **C:** A cranial window was positioned over lobules V, VI and Simplex and climbing fiber activity was imaged with two-photon microscopy after viral transfection of contralateral inferior olive neurons with GCaMP8m (top). Post-hoc immunofluorescence analysis was used to verify the expression of GCaMP (green) in climbing fibers after anti-Calbindin (red, Purkinje cells) antibody labelling (bottom). **D:** Average Ca^2+^ event probability traces (no binning, 0.03 s sampling period) of climbing fibers aligned to go cue and reward (N=954 cells, 4 mice). The fibers exhibited either “pure selectivity” (left, unimodal responses) or “mixed selectivity” (right, multimodal responses). The cells are grouped into functional categories (ordinate labels) and ordered within each category by increasing peak probability (top to bottom). **E:** Fraction of fibers in each functional category (Venn diagram, direction of arrows indicates an increase in cell numbers as they become less specifically activated by force) and their distribution across the imaged lobules V, VI and Simplex (fraction of the total responsive fibers in each lobule). ***: p<0.0001, n.s.: p>0.01 (χ^2^ test).

We virally expressed GCaMP8m in neurons of the inferior olive contralateral to the limb and imaged the axonal boutons and varicosities of climbing fibers in the ipsilateral cerebellar hemisphere with two-photon microscopy (Fig. 1C, Suppl. Movie 1). Post-hoc immunohistochemical analysis confirmed the expression of GCaMP in climbing fibers that intertwined around proximal dendritic branches of Calbindin positive Purkinje cells (Fig. 1C, bottom). We surveyed the lobule Simplex and adjacent paravermal lobuli V and VI known to receive forelimb afferents ^39^ and control ipsilateral forelimb movements ^40,41^. Climbing fibers reliably responded (significantly increased the probability of Ca^2+^ events compared to baseline, see Methods for details) to all three sensory events: force onset, go cue and reward (954 cells, 30 sessions, 4 mice, Fig. 1D). 40% responded in a unimodal manner (left panel, Fig. 1D) whereas the other 60% were multimodal, showing activation by two or all three stimuli (right panel, Fig. 1D), thereby exhibiting the property of “mixed selectivity” ^42^. Surprisingly, the force-only responding cells were the smallest category (8%) and the number of activated cells increased as the category became less force specific (Venn diagram, Fig, 1E). Therefore, in our experiment, an unexpected proprioceptive stimulation of the forelimb was less efficient in driving CF activity compared to a predictable auditory go cue or reward delivery. Force cells were significantly more numerous in lobule V, reward cells in lobule Simplex and go cue cells were equally distributed across the three lobules (Fig. 1E). Unimodal cells were more numerous in lobule VI compared to V and Simplex whereas the opposite distribution was observed for multimodal cells (Fig. 1E).

Because mice initiated forelimb movements shortly after the go cue, the cells responsive to go cue might also be motor related movement cells. To test for this possibility, we identified all CFs with significant increases in Ca^2+^ event probability when aligning correct trials to either the go cue or movement onset. The identified population of responsive CFs were more precisely time locked to go cue than to movement onset (Suppl. Fig. 1). Event probability sharply increased within 33 ms after go cue onset whereas the alignment to movement initiation was temporally broad and activity started increasing more than 100 ms before movement. We thus confirm that single CF activity is poorly related to movement ^27^ and that movement parameters might instead be encoded in the CF population activity ^38,43–45^. This result would furthermore suggest that previously reported CF activation in lobuli V, VI and Simplex of the mouse by self-initiated forelimb movements are related to the animal’s decision to produce a movement (i.e. an internal go cue) rather than motor signals ^28,33,34,40^. Self-initiated movements (without the presence of a sensory cue) also show a highly reduced CF activation compared to sensory-cued movements with matched kinematics ^40^.

In our task, we identified CFs with different response types (force vs. go cue vs. reward) and complexities (unimodal vs. multimodal). Because each of the three sensory events capable of triggering CF bursts could be followed by different behavioral outcomes (e.g. perturbed vs. adapted limb position) we explored in subsequent analyses whether different categories of CFs are informative of such outcomes.

### “Sensory saliency” vs. “motor organization” coding by climbing fibers

We asked whether force activated CFs respond differently when the force perturbs limb position (perturbed trials) compared to when the force is counteracted (adapted trials). Given the often probabilistic nature of CF activation, we used information theory ^10^ to quantify the predictive information on the behavioral outcome (perturbed vs. adapted) conveyed by the occurrence of CF events. Separating perturbed and adapted trials for each session (Fig. 2A) identified CFs with significant increases in information rate (*I_R_*) after force onset (Fig. 2E) as well as non-informative cells that produced the same generic response to the force irrespective of behavioral outcome (Fig. 2I). We refer to the former as a “motor organization” cell and the latter as a “sensory saliency” cell.

**Figure 2.**
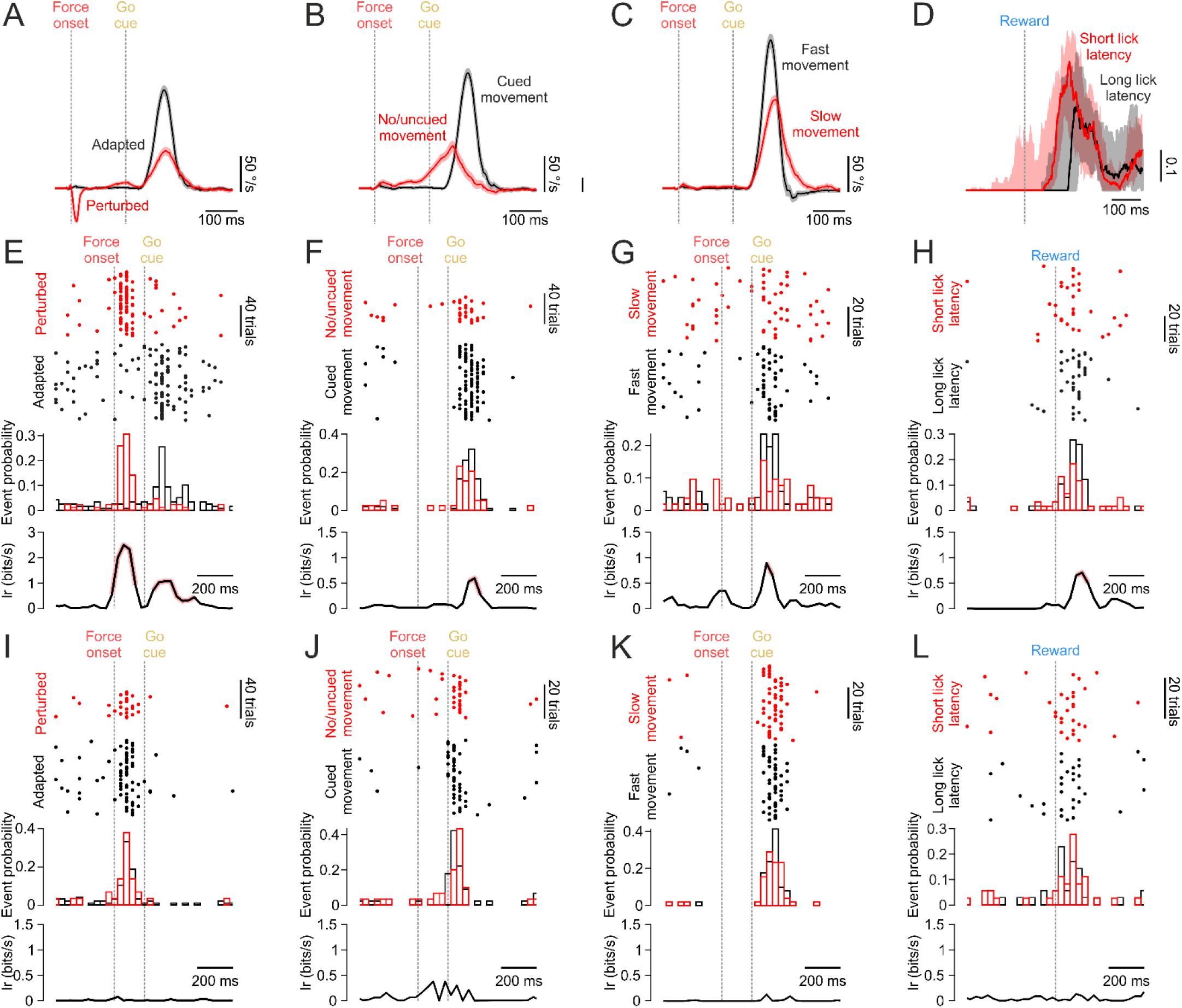
Climbing fiber activation by sensory cues can either be generic or informative about behavioral outcome. **A:** Angular velocity traces for perturbed vs. adapted trials (mean ± s.e.m. of all sessions). **B:** Angular velocity traces of adapted trials separated into cued vs. no or uncued movements. **C:** Angular velocity traces of adapted trials with cued movements separated into trials with high and low peak movement velocities. **D:** Lick probability traces in correct/rewarded trials separated into short vs. long lick latency (median ± quartiles of all sessions). **E:** Example of climbing fiber activity responsive at force onset informative about perturbed vs. adapted limb positions. Top panel: dots are detected Ca^2+^ events in perturbed (red) and adapted (black) trials. Middle panel: histograms of event probability. Bottom panel: information rate (*I_R_*) with significant values (p<0.01, χ^2^ test) highlighted in red. **F:** Same data as in E for an example fiber activated by the go cue and informative about cued vs. no/uncued movements. **G:** Example of a fiber activated by the go cue and informative about fast vs. slow movement kinematics. **H:** Example of a fiber responsive to reward and informative about short vs. long latency lick kinematics. **I-L:** same analysis as in E-H for climbing fiber examples with generic responses to the respective sensory cues.

The CF in Fig. 2E was a bimodal cell responding to both force and go cue. A significant increase in *I_R_* after go cue suggests that it is also informative about behavior in its go cue triggered activity. Because in perturbed trials the initiated movements were slow and imprecise compared to those in adapted trials (Fig. 2A, Fig. 1B) the CF might indeed carry information about these kinematic differences. Alternatively, the second *I_R_* peak (following go cue) might be a byproduct of the first peak knowing that CF bursts typically occur once or twice per second ^46^. Although occasional discharges at higher rates can be observed (up to 8 times per second) ^46^; there is an intrinsic low probability for two events to be evoked in quick succession. Accordingly, a Ca^2+^ event in response to the go cue becomes statistically less probable if the multimodal CF was triggered in the same trial by the preceding force onset. To test whether go cue activations can be classified as “sensory saliency” or “motor organization” signals we therefore only analyzed trials with adapted limb positions. The trials were separated into cued movements vs. no or uncued movements (Fig. 2B). The latter were grouped as both are instances when mice fail to respond to the go cue. We could again identify CF with significant increases in *I_R_* with respect to the behavioral outcome (Fig. 2F) as well as CF with generic activations by the go cue (Fig. 2J). We performed the same analysis on trials with cued movements only, by comparing movements with different kinematics, namely low vs. high peak velocity (Fig. 2C, G, K).

Reward triggered CF activations have been previously shown to increase their coherence at the population level (i.e. more co-activated CFs) with increased temporal precision of subsequent licking ^38^. To test whether activations of individual CFs are also informative of licking behavior we separated all correct trials into short vs. long lick latency categories and labeled cells as having either a “sensory saliency” (Fig. 2L) or “motor organization” (Fig. 2H) function.

The majority (64%, N=448) of force activated CFs could in this manner be labeled as “motor organization” cells (Fig. 3A), whereas only 17% (N=542) of go cue CFs (Fig. 3B) and 5% (N=427) of reward CFs (Fig. 3A) passed the significance threshold (p<0.01, χ^2^ test). The statistically different proportions (p<10^-91^; χ^2^ test) as well as higher *I_R_* values (Fig. 3A-C, insets; p<10^-28^, Kruskal-Wallis test) indicate that even if fewer CFs respond to a proprioceptive stimulation of the forelimb compared to go cue and reward (Fig. 1D), the former are more informative about the consequences the encoded stimulus has on behavior. The go cue activations were informative in higher proportion (p<10^-23^, χ^2^ test) about motor initiation (cued vs. no/uncued movement) than kinematics (slow vs. fast movement, 2%, N=611; short vs. long latency movement, 5%, N=611; Suppl. Fig. 2A). Interestingly, the few CFs informative about movement latency had significantly higher *I_R_* values compared to those informative about movement velocity or initiation (p<10^-7^, Kruskal-Wallis test; Suppl. Fig. 2A). Reward CFs were marginally more informative about lick latency compared to lick rate or licking precision in their proportion (p<0.01, χ^2^ test; Suppl. Fig. 2B) but not in terms of *I_R_* values (p=0.3; Kruskal-Wallis test).

**Figure 3.**
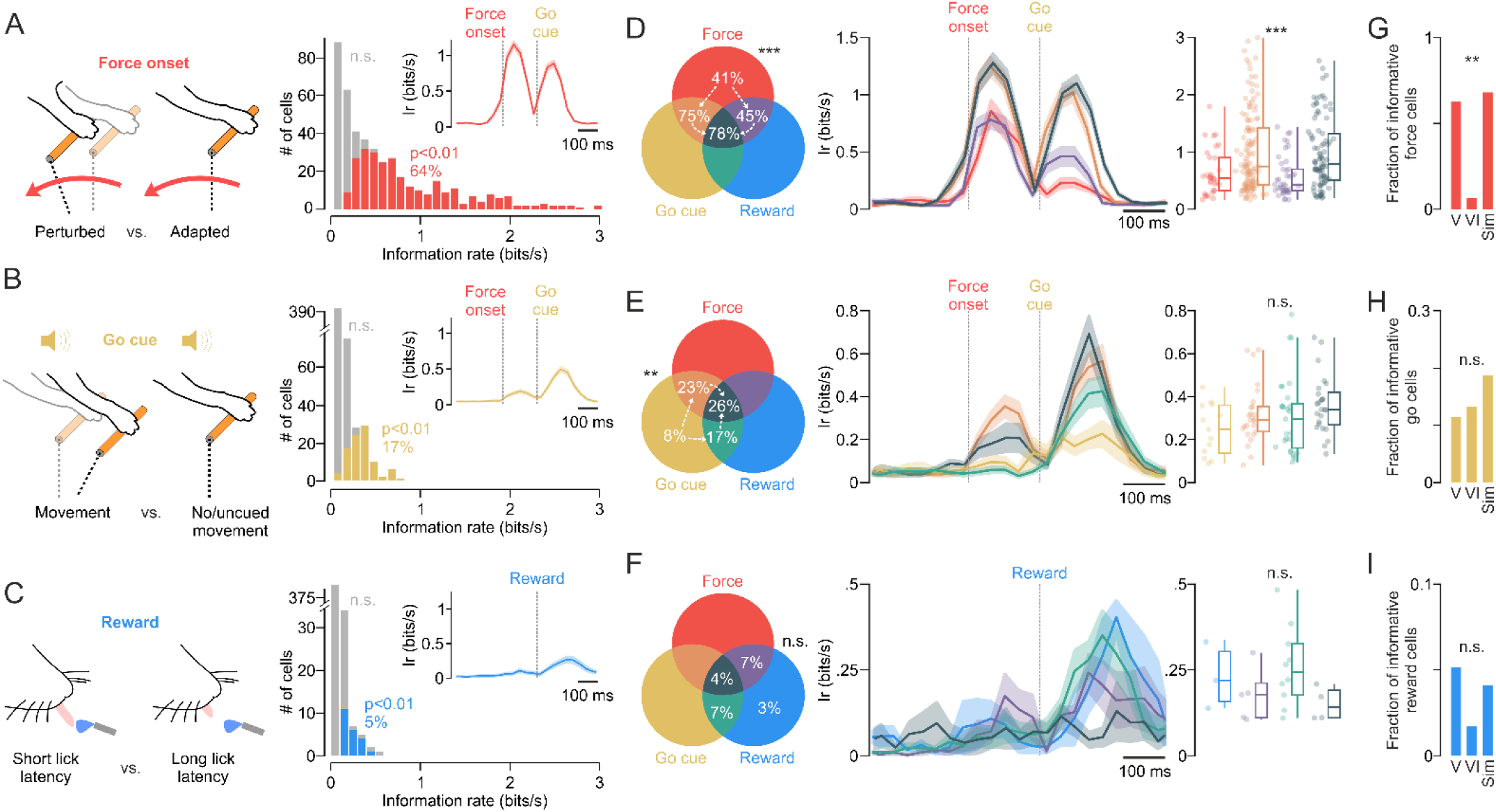
Information carried by climbing fibers is regulated by sensory event modality and the cell’s functional complexity. **A:** Distribution of information rate predictive of behavioral outcome (perturbed vs. adapted limb position) conveyed by Ca^2+^ event occurrence of all climbing fibers activated at force onset. 64% were significantly informative of behavioral outcome (red, p<0.01, χ^2^ test) and the rest had generic activations (grey). Inset: mean (± s.e.m.) information rate trace of all informative cells. **B:** Same data as in A for climbing fibers responsive to go cue that are generic (grey) or informative (yellow, 17%) of behavior (cued vs. no/uncued movement). **C:** Same data as in A,B for reward responsive fibers showing the ability of a minority of cells (5%) to convey information about lick latency. **D**, Left: Fraction of force-related informative climbing fibers in each functional category indicates that information increases for fibers with multimodal responses (direction of arrows). Middle: Information rate trace averages (± s.e.m.) for fibers with unimodal and multimodal functional properties (see Venn diagram for color code). Right: Comparison of information rate between the functional categories (box plot: median +/-quartiles and min/max values; dots: individual data points). **E-F:** Same data as in D for informative climbing fibers in B and C, respectively. **G-H:** Fraction of informative cells in each of the imaged cerebellar lobules. ***: p<10^-10^, **: p<0.001, n.s.: p>0.05 (χ^2^ test) for D-F left panels. ***: p<10^-4^, n.s.: p>0.05 (Kruskal-Wallis test) for D-F right panels. **: p<0.01, n.s.: p>0.05 (χ^2^ test) for G-I.

We next asked whether coding of behavioral information is related to the functional complexity (unimodal vs. multimodal) of CFs. For the force activated CFs, the proportion of informative cells was significantly different across the response categories (p<10^-10^, χ^2^ test). The proportion increased as the cells became more complex (Fig. 3D, Venn diagram, direction of arrows). The one bimodal (force and go cue responsive) and the trimodal categories had more than 75% of informative CFs, which were also statistically more informative (higher *I_R_*) compared to the other bimodal (force and reward responsive) and the unimodal categories (p<10^-4^, Kruskal-Wallis test; Fig. 3D). For the go cue activated CFs, the same increase in the proportion of informative cells with functional complexity was observed (p<0.001, χ^2^ test; Fig. 3E, Venn diagram). However, the information rate did not significantly differ between the functionally different categories (p=0.11, Kruskal-Wallis test; Fig. 3E) even though a post-hoc comparison between the unimodal go cue only responsive cells and the trimodal cells did yield a marginally significant difference in *I_R_* values (p=0.02, Wilcoxon rank sum test). For the few reward activated CFs informative about lick latency, no significant differences between categories were observed (Fig.3F; p=0.5, χ^2^ test; p=0.2, Kruskal-Wallis test).

In terms of topography, the fraction of CFs informative about how licking or limb reaching is organized in response to reward or go cue, respectively, did not differ in lobules V, VI and Simplex (Fig. 3H, I). CFs projecting to lobules V and simplex activated by the force stimulus were informative about the ensuing motor response in more than 66% of cells, whereas less than 6% of fibers projecting to lobule VI carried this information (p<0.01, χ^2^ test; Fig. 3G).

In summary, CFs encoding a force acting on the forelimb differ from CFs encoding a “teleceptive” movement trigger or a reward in terms of their capacity to instruct the cerebellum about the ensuing motor response. When responding to force, the CFs serving as “motor organization” cells are more numerous, produce more informative activity and are topographically more specific. The information is maximized in the CF subpopulation with multimodal functional properties, demonstrating that “mixed selectivity” neurons have higher computational flexibility compared to “pure selectivity” neurons ^42^.

### Climbing fibers encode behavioral outcomes by modulating the burst size

CFs transmit bursts containing between 1 and 6 action potentials to evoke complex spikes in the post-synaptic Purkinje cells ^47,48^. Whether CF burst size has a particular significance or is random, is a central question about the way the olivocerebellar system encodes information ^49^. In addition to modulating event probability, do CFs transmit information about the significance of sensory cues by modulating the number of spikes in individual bursts? To answer this question, we took the size of imaged Ca^2+^ events as a proxy for the number of spikes in CF bursts ^50^. We first compared events evoked by the force, go cue and reward stimuli to spontaneous events in preceding baseline periods. Evoked events had larger sizes than spontaneous events for all three stimuli (Fig. 4; p<10^-33^, t-test). The difference was however higher for the go cue compared to the force and reward responsive CFs (p<0.001, one-way ANOVA). At the single cell level of analysis, a higher fraction (p<10^-12^, χ^2^ test) of go cue (41%, N=358) than force (25%, N=175) or reward (12%, N=215) activated CFs had significantly stronger evoked than spontaneous bursts (Fig. 4, open blue symbols; p<0.01, t-test).

**Figure 4.**
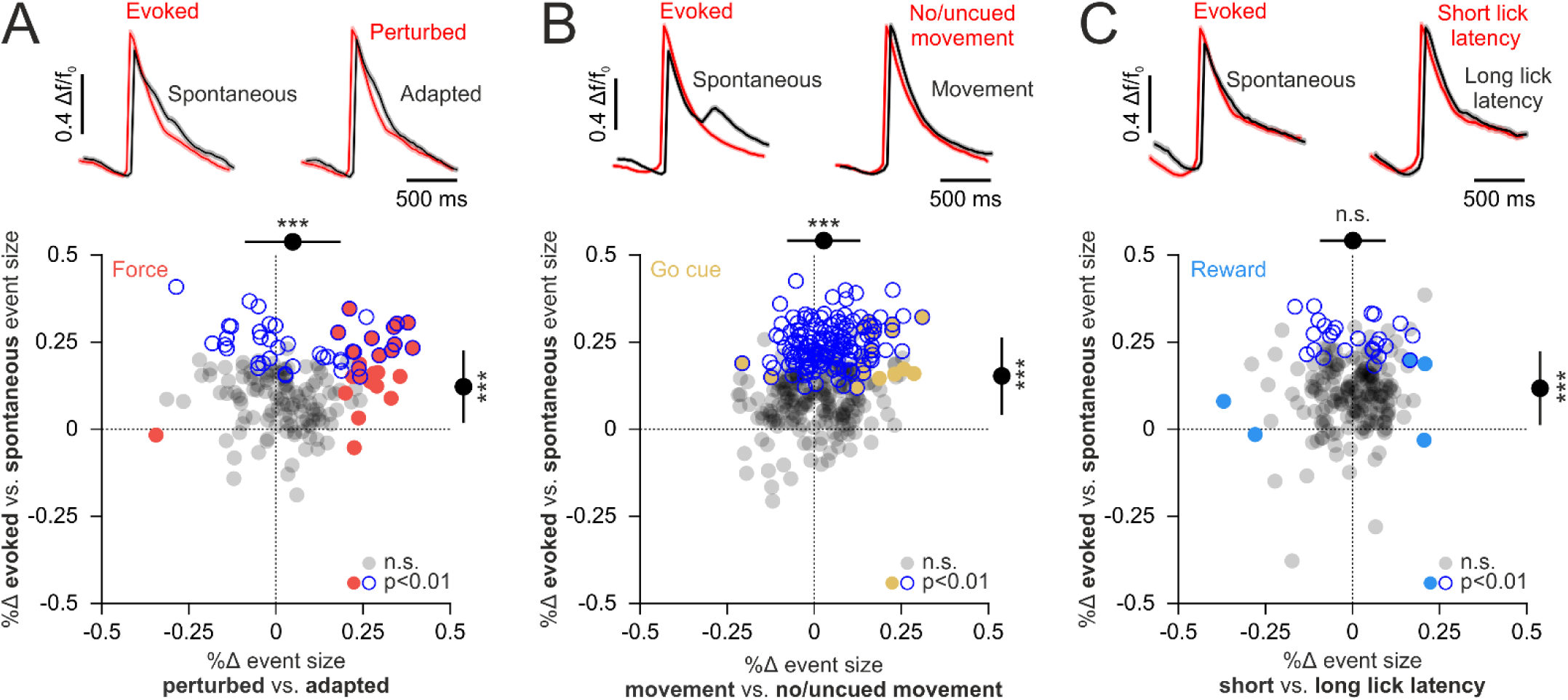
Coding of sensory events and behavioral outcomes by the size of climbing fiber Ca^2+^ events. **A**, Top: Mean (± s.e.m.) Ca^2+^ event Δf/f_0_ traces of force-related climbing fibers produced in baseline periods (spontaneous) and after stimulus onset (evoked). The evoked event Δf/f_0_ traces are further separated for perturbed vs. adapted trials. The traces are temporally misaligned for clarity. Bottom: % change (Δ) of evoked vs. spontaneous event sizes plotted against the same measure of evoked events in perturbed vs. adapted trials. Data points are individual fibers responsive at force onset. Blue open symbols: significant difference for evoked vs. spontaneous. Red symbols: significant difference for perturbed vs. adapted. Grey symbols: no significant difference in either comparison (t-test). Black filled symbols: % change means ± s.d., ***: p<10^-4^, n.s.: p>0.05 (t-test). **B, C:** Same data as in A for go cue and reward related climbing fiber events.

We next compared the size of sensory evoked CF events for different behavioral outcomes. Force and go cue activated CFs produced more spikes for adapted vs. perturbed (p<10^-4^, t-test) and cued vs. no/uncued (p<10^-5^) movements, respectively (Fig. 4). Lick latency however did not modulate the burst size of CFs responsive to reward (p=0.87, t-test). The number of cells with significant burst size differences (Fig. 4, filled symbols; p<0.01, t-test) were more numerous (p<10^-5^, χ^2^ test) for the force (15%, N=175) compared to go cue (5%, N=258) and reward stimuli (2%, N=215).

Our results therefore confirm previous reports that the CF burst size is not a random process underlying a binary “all-or-nothing” signal ^25,51,52^. The number of spikes can be used to discern spontaneous from sensory evoked CF bursts but also to transmit information about the subsequent motor response. Additionally, the strength of burst size modulation depends on the sensory modality or behavioral significance of the encoded sensory event.

### Climbing fiber activation is modulated by behavioral context

CFs activated by force were the most informative (Fig. 3A,D; Fig. 4) about the consequence on behavior (adapted vs. perturbed limb position). The hypothesized “motor organization” signal might mainly parametrize the underlying motor variables. CFs activated by reward, on the other hand, were the least informative (Fig. 3C,F; Fig. 4) about the ensuing motor response (i.e. lick kinematics). They thus mostly carry a “sensory salience” signal that might organize, not movement parameters, but a transition between behavioral states, from reward anticipation to reward consumption ^33^. Given these differences, we reasoned that the preceding wait-for-force and wait-for-reward, seemingly identical, “silent” periods (Fig. 1B) differ in terms of behavioral context. Because trials were aborted upon manipulandum release, mice had to continue holding the manipulandum during both periods. Reward and force anticipation are therefore not passive moments of inaction in our task; the mice are actively generating inconspicuous motor signals that might fundamentally differ in the two contexts. We therefore asked: does behavioral context modulate the probabilistic activity of CFs during which mice produce subliminal actions?

To answer this question, we quantified the predictive information present in CF event occurrence on period identity (wait-for-force vs. wait-for-reward; Fig. 5A). We restricted the analysis to the last 0.5 s of each period. We identified CFs with both significantly lower (Fig. 5B; reward cell; *I_R_*=0.06 bits/s; p<10^-6^, χ^2^ test) and higher (Fig. 5C; go cue cell; *I_R_*=0.08 bits/s; p<10^-7^, χ^2^ test) event probability during reward anticipation. A large number of CFs were in either of these two manners informative (37%, N=952; p<0.01, χ^2^ test) about whether the mice were waiting for the force or the reward (Fig. 5D). Interestingly, the proportion of informative cells increased as their functional category became more reward specific (Fig. 5E, Venn diagram, direction of arrows). The information rate also significantly differed between the categories (p<10^-5^, Kruskal-Wallis test; Fig. 5E), with the unimodal go cue, the unimodal reward and the bimodal go cue and reward CFs having the highest *I_R_* values. The topographic distribution of the informative cells was uniform across the three imaged cerebellar lobules (Fig. 5F; p=0.87, χ^2^ test).

**Figure 5.**
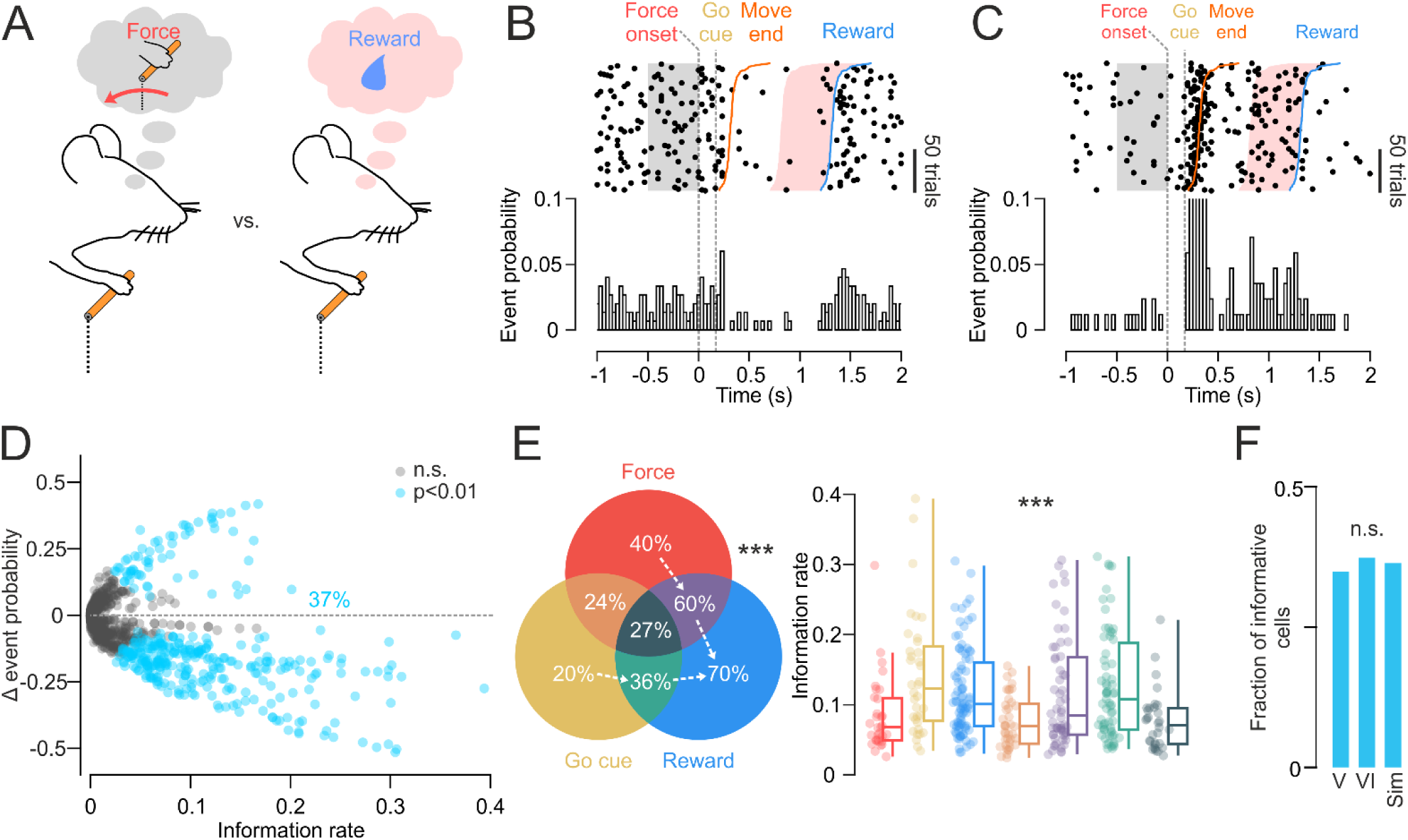
Climbing fibers encode behavioral context. **A:** comparison of climbing fiber activation during silent periods of waiting within different behavioral contexts: waiting in anticipation of force vs. waiting in anticipation of reward. **B-C:** Examples of climbing fiber activity, from the same session, informative about behavioral context with higher event probability during “wait for force” or during “wait for reward”, respectively. **D:** Difference (Δ) in event probability between the “wait for force” and “wait for reward” periods plotted against the information rate for N=952 fibers (cyan symbols: 37% of informative cells, p<0.01, χ^2^ test). **E,** Left: the fraction of informative cells increases as the responses become more reward-specific (direction of arrows, ***: p<10^-23^, χ^2^ test). Right: information rate of climbing fibers in each functional category (box plot: median +/-quartiles and min/max values; dots: individual data points, ***: p<10^-5^, Kruskal-Wallis test) **F:** Fraction of informative responses per imaged lobule (n.s.: p>0.05, χ^2^ test).

We conclude that CFs do not only encode sensory events but also behavioral context. Specifically, the probabilistic activation of CFs was modulated in our task by the nature of the anticipated sensory event (force vs. reward) during otherwise identical, seemingly idle, periods. The remaining challenge is to identify and measure the key parameters that differentiate the two contexts. The probability of manipulandum release was not significantly different in the two periods (Suppl. Fig. 3) making distinct levels of impulsivity ^53^ an unlikely explanation. Alternatively, CF activation might reflect different forms of preparatory motor activity (force counteraction vs. lick initiation) or the cost associated with aborting the ongoing behavior. Indeed, releasing the manipulandum during wait-for-force results in a shorter delay until the next reward compared to releases during wait-for-reward, thus incurring a higher cost to the latter.

### Dissociating climbing fiber from Purkinje cell dendritic signaling

Most of our knowledge about the olivocerebellar system comes from recording complex spikes or imaging dendritic Ca^2+^ signals from Purkinje cells (PC) rather than direct recording or imaging of climbing fibers. The post-synaptic PC dendritic Ca^2+^ signal or the somatic PC complex spike are assumed to be reliable representations of CF activity. Although the CF-PC synaptic transmission is highly reliable ^25,52^, the ensuing dendritic and somatic PC signals can be modulated by a number of factors ^49,54^: parallel fiber contribution to the Ca^2+^ signal ^55^, regulation of PC membrane potential by inhibitory synapses from molecular layer interneurons ^56^, inhomogeneous excitability of dendritic branches ^57,58^, aminergic modulation of synaptic excitation onto PCs ^59^, direct interaction of microglia with PC dendrites ^60^, possible innervation of a PC by more than a single CF ^61^ and plasticity-related modification of the CF-PC synapse ^62^. In our experiments, we have therefore chosen to directly image Ca^2+^ activity of CF axonal terminals in the cerebellar cortex, instead of indirectly inferring the activity from proxy signals in PCs.

In a final experiment, we asked whether the post-synaptic CF responses in PCs are, in part, graded independently of variations in pre-synaptic CF bursts. We performed simultaneous in-vivo two-photon Ca2+ imaging of CF axonal terminals and PC dendrites, after viral transfection of the contralateral inferior olive excitatory neurons with GCaMP8m, and sparse expression of cre-dependent RCaMP1.07 in lobules V, VI and Simplex of Pcp2-cre mice (Fig. 6A). As previously reported ^52^, analysis of Ca^2+^ dependent activity from identified CF-PC pairs revealed high signal correlations (r^2^=0.71±0.15, mean±s.d.; N=36 pairs, all p≈0) and reliable synaptic transmission (≈100%) quantified as the percentage of concurrent Ca^2+^ events (i.e. spike bursts) (Supplementary Movie 2). We next analyzed the relation between amplitudes of synchronous events in CF-PC pairs. We hypothesized that if the pairs can be modeled as an idealized single input, single output linear system then deviations from unity correlation should only be attributable to noise inherent to in-vivo imaging. In that case, the distribution of errors between standardized scores of CF-PC amplitude pairs should have zero mean and thus not show any residual contribution from non-CF signals. We visualized the error distribution in terms of the directly proportional distribution of distances to the unity line of the standardized score pairs (Fig. 6B) and observed that in 22/36 pairs the mean significantly deviated from zero (p<0.01, t-test). Furthermore, in 16/36 pairs, the standardized event amplitudes had different underlying distributions (p<0.01, two sample Kolmogorov-Smirnov test) suggesting that the two signals are, in part, graded with independent statistics. For comparison, we performed the same analysis between bouton/axon segments of the same CF (Fig. 6C,D) after viral transfection of the inferior olive only. As for PC-CF pairs, CF segment pairs showed both high event synchronicity (≈100%) and significant signal correlations (r^2^=0.89±0.07, mean±s.d.; N=41 pairs, all p≈0). However, the error distributions between standardized amplitude pairs had non-zero mean in 5/41 (p<0.01, t-test) and different underlying distributions in 0/41 pairs (p<0.01, two sample Kolmogorov-Smirnov test). Both ratios were significantly lower in comparison with PC-CF pairs (p<10^-5^, χ^2^ test).

**Figure 6.**
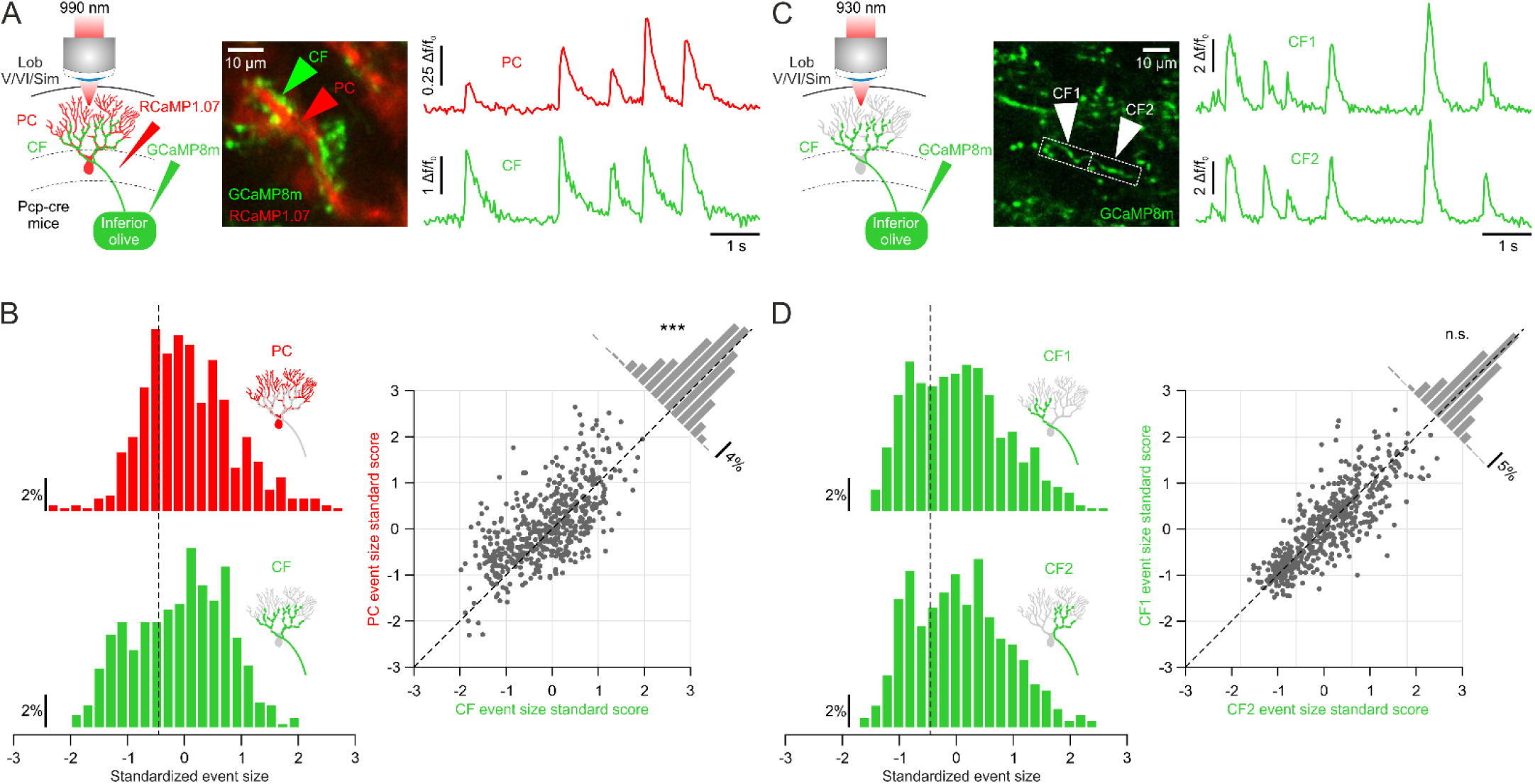
Simultaneous imaging of pre-synaptic climbing fiber and post-synaptic Purkinje cell dendritic Ca^2+^ activity. **A:** Example of dual color imaging of a climbing fiber (CF)-Purking cell (PC) pair and their Ca^2+^ dependent Δf/f_0_ fluorescence traces. **B:** The distribution of standardized event sizes was statistically different for the pair in A (left, p<0.001, two sample Kolmogorov-Smirnov test) and the mean of the errors between the standardized scores significantly deviated from zero (right, ***: p<10^-10^, t-test). **C:** Example of an identified climbing fiber segment pair (CF1 and CF2) and their respective Ca^2+^ dependent Δf/f_0_ fluorescence traces. **D:** Standardized event size distributions did not differ between CF1 and CF2 in C (left, p>0.05, two sample Kolmogorov-Smirnov test) and the mean of errors between their standardized scores was not significantly different from zero (p>0.05, t-test).

In summary, it seems that PC dendrites integrate graded CF bursts into graded post-synaptic responses ^25^ but not in a one-to-one linear manner. The non-linearity suggests that regulation of PC Ca^2+^ dendritic signaling by factors other than the CF input does indeed occur in-vivo. On the other hand, the disparities between event amplitudes of a few CF segment pairs likely reflect decreased signal to noise ratio due to the small size of the imaged structures. Alternatively, differential modulation of boutons of the same CF might be possible via signaling in the extracellular space ^63^, but not in the form of axo-axonic synapses ^64^, from molecular layer interneurons.

## Discussion

The role of the cerebellum is to ensure reliable motor responses to stimuli. From that perspective, PCs act as a hub for selecting incoming sensory and motor signals forwarding only what is important for coordinating behavior to the cerebellar output nuclei for final processing. The single climbing fiber input to a PC seems to play a major role in this selection process. It regulates the efficacy of tens of thousands of parallel fiber synapses^65–70^ that relay a barrage of sensorimotor information from the mossy fiber inputs. In this study, we analyzed to what extent the influence of CF activation is observed in the behavioral response, meaning how likely they are to be drivers of “motor organization”. Our findings show that this function is more attributable to CFs that encode bottom-up inputs from the somatosensory periphery than to those encoding top-down cortical instructions. Additionally, it is more prevalent in CFs with multimodal selectivity and can be operative in the absence of sensory stimulation.

### Climbing fibers as versatile signal encoders

CFs can be activated by a broad range of stimuli ^25,26,28,31,38^ but seem more sensitive to somatosensory than to “teleceptive” auditory or visual signals ^26,27^. This is consistent with the underlying anatomical pathways: somatosensory afferents are directly passed to the inferior olive via second order neurons in the spino-olivary tract ^71–73^, whereas auditory or visual signals may be conveyed indirectly through higher-order structures such as the mesodiencephalic junction ^35,36^ and the superior colliculus ^74^.

We therefore expected CFs to predominantly respond to force onset but found that the majority were activated by the go cue and reward (Fig. 1D). We propose that CFs do not just encode sensory input but adapt to relevant task cues through learning. In our experiment, the go cue signals the need for action (i.e. is a movement instruction) while force onset requires no explicit response or adaptation for correct task execution. Because we started imaging only after the task was learned, we suspect that more CFs were responsive to the somatosensory stimulation in naïve animals and gradually adopted task relevant responses as learning unfolded and the go cue and reward gained behavioral significance. This refinement gives salient stimuli meaning and reduces responses to less important stimuli ^32,37^, contrasting with responses observed when animals are passively exposed to stimuli without a structured task ^25–27^.

Most CFs had multimodal responses, responding most often to two or even all three sensory cues (Fig. 1D). Our task unambiguously highlights their multimodal nature because the stimulus sequence was learned, and each had a specific behavioral relevance. The activations thus cannot simply be described as generic responses to unexpected sensory events as in previous studies ^25,26^. The experiment occurred in a controlled environment involving a few sensorimotor variables. We therefore expect CFs to exhibit an even higher degree of mixed selectivity when mice act in real-world conditions. It is not surprising that mixed selectivity is also a feature of climbing fibers given that inferior olive neurons receive highly convergent inputs and are coupled by gap junctions ^75,76^. The property of mixed selectivity ^42^ suggest that circuit motifs underlying functions such as adapting to a perturbation, initiating cued movements or processing reward are overlapping in the olivocerebellar system, which is thereby endowed with the necessary flexibility to organize diverse and complex forms of behavior. Despite such overlap, the different representations still maintain a degree of topographic specificity in their projections to the cerebellar cortex (Fig. 1E).

### Climbing fibers as informative organizers of behavior

Our results show a stark contrast between the information about perturbed vs. adapted limb positions conveyed by force activated CFs, and the information that go cue or reward responsive CFs contain about movement initiation or its kinematics (Fig. 3,4). Why are CFs more informative about an implicit adaptive behavior compared to explicit motor actions? One answer is that implicit motor adaptation is hardcoded in the spino-olivocerebellar circuit (i.e. is an archaic function) whereas organizing voluntary actions in response to instructive cues requires associative learning based on top-down cortical inputs to the inferior olive (i.e. a newer cerebellar function) and is hence underrepresented. Following the reasoning in the previous paragraph, the difference might also reflect an ongoing temporal evolution of CF encoding. CFs might first have to be calibrated towards responding to a sensory cue, and only after calibration does their activity start being predictive of and orchestrating the motor response ^77^. As our imaging results provide only a snapshot of CF activity, we hypothesize that over time, go cue and reward activations would show increased information about the influence the encoded signals exert on behavior.

An intriguing finding was that multimodal CFs were more likely to be informative about behavior and more strongly encode this information (Fig. 3D, E). This suggests that mixed selectivity CFs do not only confer flexibility about the multiple sensory signals they encode but also about the impact each of these signals has on behavior. In other words, CFs selective for multiple signals in one dimension (e.g. sensory stimuli) are also selective for multiple signals in another dimension (e.g. motor actions or context), whereas a pure selectivity CF representing one stimulus is also likely to lack flexibility across other dimensions. This functional dichotomy highlights the unique role of multimodal neurons, not merely as multitaskers but as key players in integrating and adapting to behavioral demands.

### Representation of behavioral context

CFs are spontaneously active in the absence of external stimuli during seemingly idle periods, raising questions about the nature and purpose of such “noise” activations ^49^. We compared CF activation in two such idle periods and found that it is statistically different when mice wait in anticipation of force onset or in anticipation of reward in 37% of cells (Fig. 5). We argue that such activations are not “noise” and instead encode specifics of the behavioral context. Similar, but overtly different, modulations of PC complex spike activity have recently been reported ^31,78^. They manifested as either buildup activity before a motor response in a small subset of cells ^78^ or brief transient increase following trial onset ^31^. These are fundamentally different from the bidirectional differences of sustained CF activity between the two periods observed in our experiments (Fig. 5B-D).

The differential activation of CFs might reflect putative olivary inputs originating in premotor cortical areas providing distinct preparatory motor signals in the two contexts. Although expectation or preparatory motor activity is observed in the cerebellum ^78–81^ it has to our knowledge only been attributed to the cortico-cerebellar loop implicating the mossy fiber system. Indeed, preparatory activity prior to licking is observed in PC simple spikes and not complex spikes ^78^, suggesting that the olivocerebellar system contributes little to motor preparation. Moreover, we found no topographic preference for the CFs informative about behavioral context (Fig. 5F). It therefore does not reflect the topographical preference of either force or reward responsive CFs (Fig. 1E), which would be expected if the modulation reflected preparatory forelimb vs. licking motor signals.

Releases resulted in aborted trials during “wait for reward” and delayed trial start during “wait for force”. Mice were therefore engaged in active holding of the manipulandum in both waiting periods. Serotonergic signaling from the raphe nucleus has been suggested to support the ability to wait, either by suppressing impulsivity ^82^ or promoting persistence ^53^. Optogenetic activation of serotonin neurons increases the animal’s ability to wait for delayed rewards ^83,84^ and their sustained activity during water/food delays ^85^ resembles the CF modulations we observed in the “wait for reward” period (Fig. 5B). Our findings might thus reflect the modulation of the inferior olive by serotonergic afferents ^86–91^ even though our crude measure of impulsivity did not show a difference between the “wait for reward” and “wait for hold” periods (Supplementary Fig. 3).

Activation of dorsal raphe serotonin promotes patience but is itself not rewarding ^83^. We however found that CFs informative about the two waiting contexts were most likely to be responsive to reward (Fig. 5E). This raises the alternative possibility that dopaminergic modulation of the inferior olive ^92–94^ might underlie the observed CF signaling. Dopamine neurons of ventral midbrain also exhibit sustained activity in anticipation of reward and seem to encode uncertainty about motivationally relevant stimuli ^95^. In our task, reward is motivationally more relevant than the force stimulus and the possibility of aborting the trial due to release makes the reward uncertain, thereby supporting the dopaminergic modulation hypothesis.

Regardless of the underlying neural circuit mechanisms, our findings highlight the versatile role of climbing fibers in encoding not just transient sensorimotor events but also sustained context-dependent signals. We conclude that CFs are highly flexible encoders of possibly any process that shapes behavior.

## Materials and Methods

### Mice

For climbing fiber imaging and behavioral experiments we used C57BL/6 mice (Janvier Labs; 2 males, 2 females, 8 to 12 weeks of age at the start of experiments). For dual imaging of Purkinje cells and climbing fibers, we used Pcp-cre mice (2 females, Erasmus University, 10 weeks of age). Mice were housed in an animal facility in groups of up to five mice per cage, maintained on a 12h/12h light/dark cycle and placed on a water restriction regime of 1 ml/day during experiments. All procedures were approved by and complied with the guidelines of the Fribourg Cantonal Commission for Animal Experimentation.

### Surgery

All surgeries were performed under 1.5% isoflurane anaesthesia. We administered additional analgesic (0.1 mg/kg buprenorphine intramuscular (i.m.)), local anaesthetic (75 µl 1% lidocaine subcutaneous (s.c.) under the scalp), and anti-inflammatory drugs (2.5 mg/kg dexamethasone i.m. and 5 mg/kg carprofen s.c.). Mice were fixed in a stereotaxic frame and placed on a heating pad (37°C). An incision was made over the midline between the ears and eyes to expose the scalp. To access the inferior olive, neck muscle tendons were detached. A small incision was made in the dura at the foramen magnum allowing the insertion of a bevelled injection pipette (Wiretroll II, Drummond Scientific) into the brainstem at an angle of 45° (dorso-ventral). We targeted the medial and dorsal accessory inferior olive and performed six viral injections in total (AP: −7.1 mm, ML: 0.15 mm, DV: 6.1 mm; AP: −7.25 mm, ML: 0.15 mm, DV: 5.95 mm; AP: −7.1 mm, ML: 0.5 mm, DV: 6.1 mm; AP: −7.25 mm, ML: 0.5 mm, DV: 5.95 mm; AP: −7.1 mm, ML: 0.7 mm, DV: 6.1 mm; AP: −7.25 mm, ML: 0.7 mm, DV: 5.95 mm) of AAV1-CamKII-GCaMP8m (100 nL, 1:2 dilution, 7.4 x 10¹² vg/ml titer, Zurich Viral Vector Facility). For Purkinje cell imaging, we performed 4 injections across lobules V/VI and Simplex of the right cerebellar hemisphere at 3 depths per injection targeting the PC layer (250 µm, 300 µm and 350 µm) of AAV1-hSyn-dlox-RCAMP1.07 (25 nL, 1:10 dilution, 8.9×10^12^ vg/ml titer, Zurich Viral Vector Facility). Injections were delivered over 10 min at a rate of 10 nL/min. After each injection, the pipette was held in position for an additional 5 to 10 min before moving to the next site. After injections, the neck muscle tendons were glued using cyanoacrylate adhesive (Loctite 401) to the skull slightly below their original locations. A custom-made stainless steel head bar was glued and cemented to the skull for head fixation during experiments using dental cement (Paladur, Kulzer GmbH). A circular craniotomy (1.8 mm diameter) was then performed over lobules V/VI and Simplex of the right cerebellar hemisphere. Dexamethasone was applied to the brain surface after removing the bone and a double glass window (1.8 mm lower glass and 2 mm upper glass diameter) was placed on the cerebellum and glued/cemented to the skull. The two glass layers were 150 µm thick each and bonded with optical adhesive (NOA 61, Norland).

### Forelimb movement task

#### Robotic manipulandum

We used a custom-built robotic manipulandum consisting of a single-arm linkage mounted on a DC motor (DCX22L EB SL 9V, Maxon) with integrated optical rotary encoder (ENX 16 RIO, 32768 counts/turn, Maxon). A handle with 1.85 mm diameter and 9 mm length was mounted at the endpoint for holding. The handle was connected to a capacitive sensor (MPR121, Adafruit) interfaced with a microcontroller (Arduino Nano) allowing the detection of holding and releasing. The motor was operated in position and current modes using a positioning PID controller (EPOS 24/2, Maxon) at 1 kHz sampling rate and interfaced with Matlab using EPOS2 libraries. The angular position and velocity of the manipulandum were recorded and monitored online at a 1 kHz sampling rate with a custom circuit. The quadrature signals from the optical encoders were decoded using the hardware quad decoders of Arduino DUE and the 16-bit digital signals converted to analog signals with a DAC (AD669ANZ, Analog devices) and provided as inputs to the Analog Input Module of the Bpod system (Sanworks).

#### Behavioral training

We used the Bpod State Machine r1 system (Sanworks) interfaced with Matlab to control and measure behavioral output. We created a Matlab object plugin to control the robotic manipulandum from within a Bpod protocol.

7 days after surgery, mice were placed on a 1 ml/day water restriction regime and gradually habituated to head fixation and the experimental setup. Mice sat in a 3D printed tube and head fixated at a 30° forward pitch angle. The training started with a passive task in which mice held the manipulandum for 2 s followed by a passive angular displacement that progressively increased from 2° to 15° (2 mm to 11 mm Euclidean distance). Movement onset was signaled with an auditory cue (30 ms, 3.75 KHz tone located behind the mouse). After the target was reached, mice had to continue holding for 1 s to receive a reward (water droplet delivered with a miniature solenoid valve, LHDA1231215H, The Lee Company) signaled by an auditory tone (300 ms, 2.625 kHz, located in front of the mouse). Following 7 to 12 days of passive training, the protocol switched to the active task. After 2 s of continuous holding, the manipulandum switched to current control mode and applied a resistive force (progressively increased from 0.087 mNm to 2.03 mNm) followed by the go cue 170 ms later. Mice had to actively push the manipulandum and reach the target (varied between 12° and 16°) within 2 s. Upon target reach, the manipulandum was switched to position mode and followed by reward delivery as in the passive task. A new trial started once the mice stopped licking for at least 1 s. The 2 s baseline holding period was reset upon detected releases. Any release thereafter, failure to reach the target or premature movements (reaching the target prior to go cue) resulted in an aborted trial and a 2 s timeout (5 s for the passive training). Neuronal imaging started once mice reached at least 50% correct performance. The force was adjusted individually for each mouse and session and varied between 0.79 mNm and 2.03 mNm to balance the number of perturbed and adapted trials.

### Two-photon microscopy

Ca^2+^ imaging in the mouse cerebellar cortex was performed with a custom built two-photon microscope based on an open-source design (MIMMS 2.0, janelia.org/open-science) and controlled with Scanimage 5.7_R1 software (Vidrio Technologies) and National Instrument acquisition hardware. The Ti:Sapphire excitation laser (Tiberius, Thorlabs) was tuned to 930 nm for GCaMP imaging and 990 nm for dual GCaMP and RCaMP imaging. The laser was focused with a 16x 0.8 NA objective (Nikon) below the cerebellar surface, modulated with Pockels cell (350-80-LA-02, Conoptics) and calibrated with a Ge photodetector (DET50B2, Thorlabs). A 315 × 315 µm area of the cortex was scanned for climbing fiber imaging and a 135 × 135 µm area for dual color imaging, with a frame rate of approximately 30 frames/s using a resonant-galvo scanning system (CRS8K/6215H, CRS/671-HP Cambridge Technologies). Emitted fluorescence was detected with GaAsP photomultiplier tubes (PMT2101, Thorlabs) and the acquired 512 x 512 pixel images were written to disk in 16 bit format. For dual color imaging, the emitted fluorescence was split with a beamsplitter (T 570 LPXR, Chroma) and filtered with bandpass filters (531/46 BrightLine HC and 630/92 BrightLine HC, Semrock) prior to detection by PMTs.

### Immunohistochemistry

Mice were anesthetized with pentobarbital (65 mg/kg) and transcardially perfused with 4% paraformaldehyde (PFA, in 0.1 M sodium phosphate buffer, pH 7.4). The brains were dissected and post-fixed overnight in the same solution, rinsed (3×20 min) with 0.1 M phosphate buffer saline (PBS), and on successive days incubated in 15% and 30% sucrose solutions (in PBS) at 4◦C for cryoprotection. Cryoprotected cerebellum was cut in 40 μm thick sagittal sections using a microtome (MICROM HM 440E, Microm International GmbH, Walldorf, Germany). Sections were permeabilized for 80 minutes in 0.1 M PBS containing 3% Triton X-100 and blocked for 1.5 hours at room temperature in a solution containing 0.1 M PBS, 0.05% Triton X-100, 0.05% Tween-20, 10% normal donkey serum (NDS, Jackson ImmunoResearch, catalogue #: 017-000-121), and 0.02% sodium azide. The sections were washed three times for five minutes with PBS before staining overnight at 4°C with the primary antibody (Calbindin D-28k, 1:5000, Swant, Monoclonal, catalogue #300) in 0.1 M PBS containing 5% NDS. The next day, the sections were washed three times in a wash buffer (1% NDS, 0.05% Tween-20, 0.02% sodium azide in 0.1 M PBS) and incubated overnight with the secondary antibody (Alexa Fluor 594, donkey anti-mouse, 1:800, catalogue #715-585-150, Jackson ImmunoResearch) in 0.1 M PBS containing 5% NDS, 0.05% Tween-20, and 0.02% sodium azide. Sections were washed the following day (3×20 min) in the wash buffer, followed by PBS (3 x 15 min) before mounting on Superfrost Plus Adhesion slides (Fisher Scientific AG, Reinach, Switzerland) and coverslips using vectashield (antifade mounting medium). Imaging was performed using confocal microscopy (Leica STELLARIS 8 FALCON, Leica, Germany) and analyzed with LAS X software. Post-acquisition image processing was carried out with ImageJ/Fiji software (NIH, Bethesda, MD).

### Data analysis

#### Motion correction

A custom MATLAB registration script was used to correct for movement artefacts. Each acquired image was aligned to a baseline average template image recorded at the start of each session. We computed the cross-correlation between each frame and the template by multiplying the two-dimensional discrete Fourier transform of one with the complex conjugate of the Fourier transform of the other and taking the inverse Fourier transform of the product. The location of the peak cross-correlation value gave the vertical and horizontal correction shift in pixels. 10% of each image was cropped at the boundaries before carrying out the computation.

#### Region of interest and Ca^2+^ activity generation

Using the session mean and variance images, bouton/varicosity centers of active fibers with clearly identifiable morphologies were manually initialized. Regions of interest of individual fibers and background were then identified as spatial footprints using the constrained nonnegative matrix factorization method ^96^ from the CaImAn Matlab toolbox (github.com/flatironinstitute/CaImAn-MATLAB). Synchronously active and spatially connected footprints were merged. The time-varying calcium activity of each fiber (i.e. its spatial footprint) and their time-varying baseline fluorescence was subsequently extracted from the acquired images and used to compute *Δf/f_0_* traces used for analysis.

#### Ca^2+^ event detection

Spike rate was inferred from the *Δf/f_0_* traces using the OASIS deconvolution algorithm ^97^ of the OASIS toolbox (github.com/zhoupc/OASIS_matlab) or the Suite 2P toolbox (github.com/cortex-lab/Suite2P). Detected spike bursts were used to identify the onset of climbing fiber Ca^2+^ events. The size of Ca^2+^ events was calculated as the difference in *Δf/f_0_* values between onset and peak (detected as the latest increasing value following onset). Given that event sizes were not normally distributed, the standardized scores (Fig. 6) were calculated based on median and median absolute deviation (modified Z score).

#### Detection of sensory evoked responses

Climbing fibers were deemed responsive to the force, go cue or reward based on a comparison of event probability in a stimulus period compared to a temporally matched baseline period. The cell was responsive if the fraction of trials with climbing fiber events was significantly different in the two periods (p<0.01, χ^2^ test). For the force, the stimulus period was the time between force and go cue onset and the baseline period the time-matched interval preceding force onset. For the go cue, the stimulus and baseline periods were 250 ms intervals after the go cue and prior to force, respectively. For the reward, stimulus and baseline periods were defined as 250 ms intervals after and before reward onset, respectively.

#### Information analysis

To identify climbing fibers with activity informative about a given behavioral outcome we separated trials into categories. Adapted vs. perturbed trials (Fig. 2A) were identified based on the peak angular velocity of the manipulandum in the first 50 ms after force onset. The latter typically had a bimodal distribution across trials with perturbed position having negative values and adapted positions near zero or slightly positive values. The separation threshold was selected as the value best separating the two modes of the distribution. Cued vs. no/uncued movements (Fig. 2B) were identified based on the time or absence of movement onset. Movement onset was defined as the latest time sample prior to the time of peak velocity when the velocity is below 20 °/s. Trials with movement onset prior to go cue or peak velocities below 20 °/s were classified into the no/uncued movement category. Fast vs. slow (Fig. 2C) or short vs. long latency movements were split based on the median peak velocity or movement onset latency in each session, respectively. The same criterion was used for splitting trials based on lick kinematics (Fig. 2D). Lick precision (Supplementary Figure 2) was calculated as previously described ^38^.

The predictive information (*I_CF_*) on the behavioral outcome conveyed by the occurrence of CF events was calculated as the decrease in entropy in trial category (Kitazawa et al., 1998):

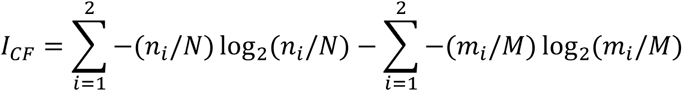

where *n_i_* and *m_i_* are the total number of trials and the number of trials with CF events in the *i^th^* behavioral outcome, respectively. *N* and *M* are the total number of trials and the number of trials with CF events in both behavioral conditions. The amount of information carried by the absence of CF (*I_CF-_*) events was calculated as:

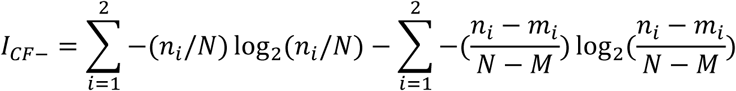

Information carried by CF events is then defined as:

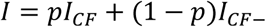

where *p* is the probability of CF event occurrence in the time interval *T* being analyzed. The information transmission rate (in bits/s) is then calculated as *I_R_* = *I/T*. For the information rate traces in Fig. 2 and 3 we used a moving window of 4 imaging frames (i.e. *T*=132 ms). In all other *I_R_* calculations, we used *T* values corresponding to analysis periods defined in the previous paragraph, except for the data in Fig. 5 where we used *T*=0.5 s. Climbing fibers were deemed informative based on the corresponding χ^2^ test and p<0.01. For all comparisons, only cells with at least 10 trials in either condition were analyzed.

#### Statistics

Measurements for any experiment were made from different animals/cells and no cells was measured repeatedly for the same analysis. Normality assumptions were made with the Kolmogorov-Smirnov test. Non-parametric tests were used when the normality assumption was not met. Significant is denoted only when p<0.01.

## Acknowledgments

We thank Carmen Schäfer and Mélanie Palacio Manzano for comments on the manuscript. This work was supported by the Swiss National Science Foundation (grant PCEFP3 181274).

## Supplementary Figures

**Supplementary Figure 1.**
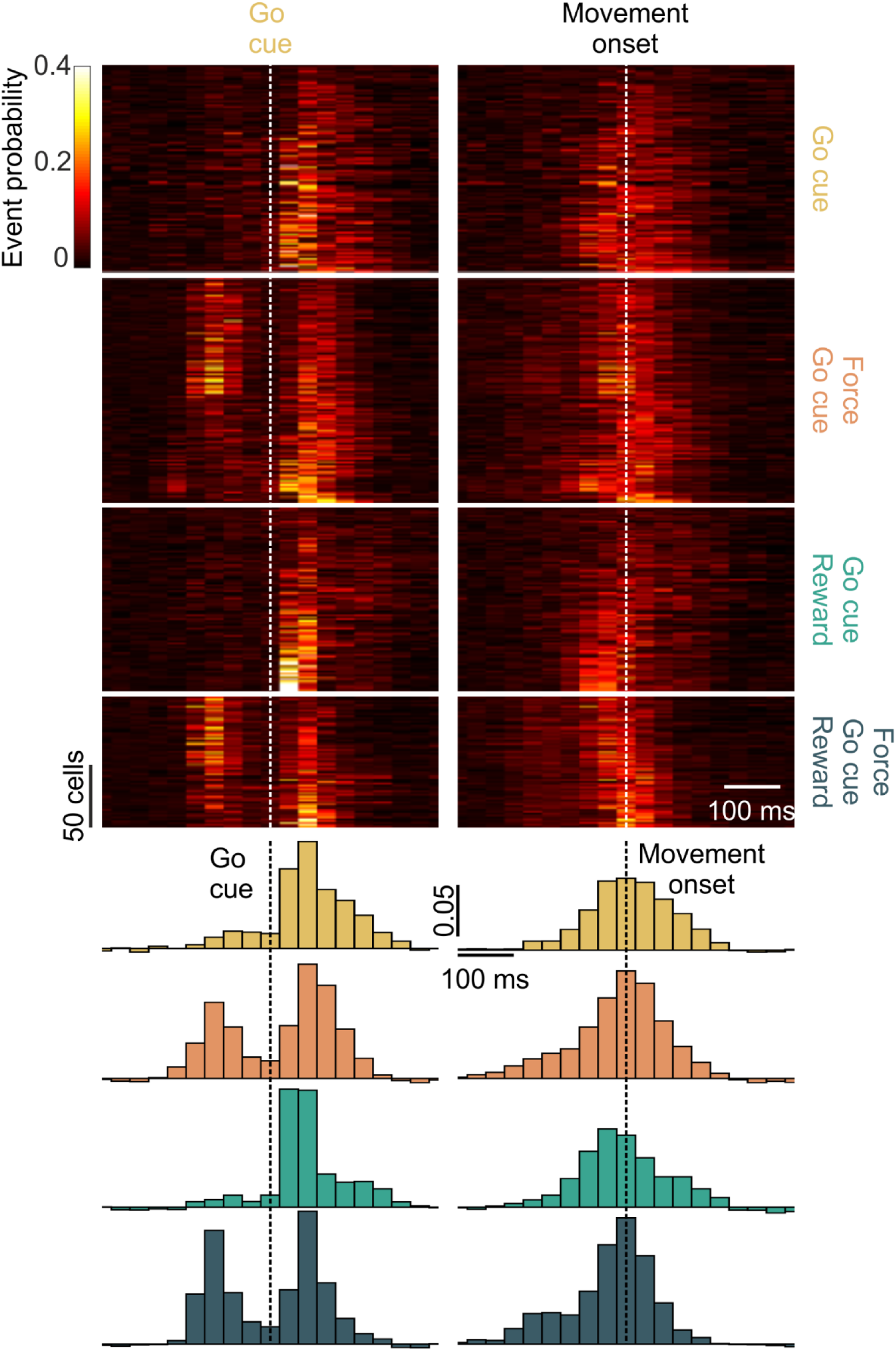
Climbing fiber activations show more precise temporal alignment with go cue than movement onset. **A:** Average Ca^2+^ event probability traces of go cue activated climbing fibers aligned to go cue (left) and movement onset (right). The cells are grouped into functional categories (ordinate labels) and ordered within each category by increasing peak probability (top to bottom). Histograms show population cell averages for the four respective functional categories aligned to go cue (left) and movement onset (right).

**Supplementary Figure 2.**
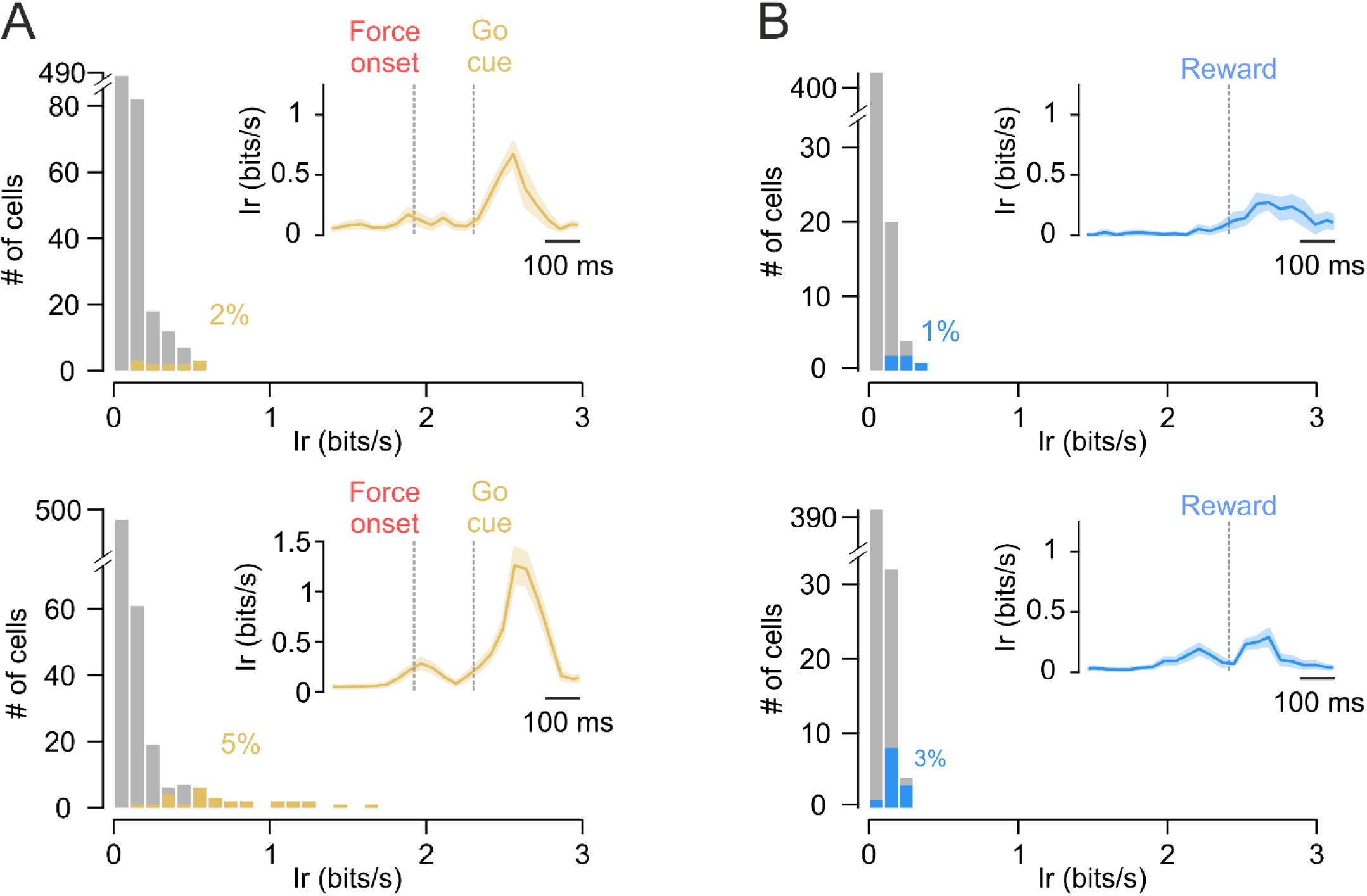
Weak modulation of climbing fiber activity by movement kinematics. **A:** Distribution of information rate predictive of behavioral outcome (Top: fast vs. slow movements. Bottom: short vs. long latency movements) conveyed by Ca^2+^ event occurrence of all climbing fibers activated at go cue. 2% (N=611) were significantly informative of movement velocity (yellow, p<0.01, χ^2^ test), 5% were informative of movement latency and the rest had generic activations (grey). Inset: mean (± s.e.m.) *I_R_* trace. **B**: Same data as in A for reward responsive fibers showing the ability of a few cells to convey information about lick rate (top, 1%, N=427) or lick precision (bottom, 3%, see Methods for details).

**Supplementary Figure 3.**
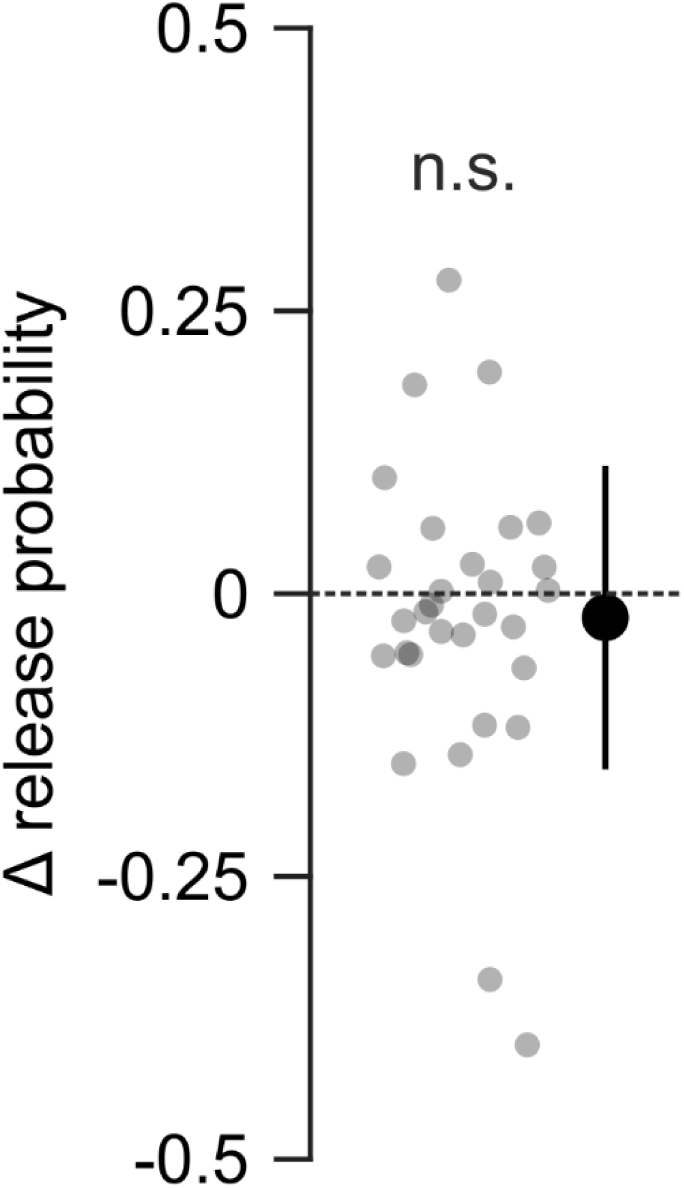
Mice were equally impulsive in anticipation of force and reward. Difference (Δ) in release probability of each session (grey points) between the “wait for force” and “wait for reward” periods. The average difference (± SD, black symbol) was not statistically different from zero (n.s.: p>0.05, t-test).

**Supplementary Movie 1. In vivo two-photon microscopy imaging of climbing fiber Ca2+ fluorescence in the mouse cerebellum.**

**Supplementary Movie 2. Simultaneous dual color imaging of pre-synaptic climbing fiber and post-synaptic Purkinje cell pairs.**

